# *Tractor*: A framework allowing for improved inclusion of admixed individuals in large-scale association studies

**DOI:** 10.1101/2020.05.17.100727

**Authors:** Elizabeth G. Atkinson, Adam X. Maihofer, Masahiro Kanai, Alicia R. Martin, Konrad J. Karczewski, Marcos L. Santoro, Jacob C. Ulirsch, Yoichiro Kamatani, Yukinori Okada, Hilary K. Finucane, Karestan C. Koenen, Caroline M. Nievergelt, Mark J. Daly, Benjamin M. Neale

## Abstract

Admixed populations are routinely excluded from medical genomic studies due to concerns over population structure. Here, we present a statistical framework and software package, *Tractor,* to facilitate the inclusion of admixed individuals in association studies by leveraging local ancestry. We test *Tractor* with simulations and empirical data focused on admixed African-European individuals. *Tractor* generates ancestryspecific effect size estimates, can boost GWAS power, and improves the resolution of association signals. Using a local ancestry aware regression model, we replicate known hits for blood lipids in admixed populations, discover novel hits missed by standard GWAS procedures, and localize signals closer to putative causal variants.

## Introduction

Admixed groups, whose genomes contain more than one ancestral population such as African American and Hispanic/Latino individuals, make up more than a third of the US populace, and the population is becoming increasingly mixed over time^1^. Many common, heritable, diseases including prostate cancer^2–5^, asthma^6–9^, and several cardiovascular disorders such as atherosclerosis^10,11^ are enriched in admixed populations of the US. However, only a minute proportion of association studies address the genetic architecture of complex traits in such groups^12,13^; admixed individuals are systematically removed from many studies due to the lack of methods and pipelines to effectively account for their ancestry such that population substructure can infiltrate analyses and bias results^14–21^. Large-scale efforts to collect genetic data alongside medically-relevant phenotypes are beginning to focus more on non-Eurasian ethnic groups that contain higher amounts of admixture^22–27^, motivating the timely development of scalable methods to allow well-calibrated statistical genomic work on these populations. If not addressed, this inability to analyze admixed people will limit the clinical utility of large-scale data-collection efforts for minorities, exacerbating the concerning health disparities that already exist^28–32^.

In GWAS, the specific concern regarding including admixed participants is obtaining false positive hits due to alleles being at different frequencies across populations. Most studies currently attempt to control for this by using Principle Components (PCs) in a linear or linear mixed model framework. However, PCs capture broader admixture fractions, and individuals’ local ancestry makeup may differ between case and control cohorts even if their global fractions are identical. Even including PCs as covariates, then, still leaves open the possibility for false positive associations, as well as absorbing power.

Studying diverse populations in gene discovery efforts not only reduces disparities but also benefits genetic analysis for individuals of all ancestries. Perhaps the most notable example of this is in multi-ethnic fine-mapping, which can dramatically reduce the variant credible set by leveraging the differing LD structures observed across populations^33–38^. This is particularly helpful in populations of African descent, where LD blocks are the shortest and individuals have nearly a million more variants per person than individuals outside of the continent^39^. We find that with admixed populations we not only can utilize the LD patterns from multiple ancestries, but have further disrupted LD blocks within each one, offering a more refined LD landscape with which to localize GWAS signal.

To help ensure that advances in genomic medicine will apply globally, we have developed a scalable framework that allows for the easy incorporation of admixed individuals into psychiatric genomics efforts by using local ancestry inference (LAI). Our framework, distributed as a scalable software package named *Tractor,* generates ancestry dosages at each site from input LAI calls, extracts painted haplotype segments for correction at the genotype level and runs a local ancestry-aware regression model, producing ancestry-specific effect size estimates and *p* values. Through testing in simulations and with empirical data on phenotypes with differing levels of polygenicity, we demonstrate that *Tractor* produces accurate results in admixed cohorts and boosts GWAS power across many genetic contexts. We further demonstrate improvements in association signal localization from the higher resolution of haplotype breakpoints in admixed genomes. These efforts fill a gap in existing resources and will improve our understanding of complex diseases across diverse populations.

The incorporation of local ancestry into variant identification for admixed populations is a concept that has been discussed previously^40–50^, particularly with regard to ‘admixture mapping,’ whereby researchers associate an elevation of a given ancestry at a locus in the genome with increased risk of a disease that is known to be stratified in prevalence across ancestries^51–55^. Admixture mapping has proven successful in diseases which are highly stratified across populations, such as asthma and cardiovascular phenotypes^56–61^. We build upon this important work by modeling the local ancestry haplotype dosage for each person at each variant in a way that allows for the generation of ancestry-specific effect size estimates while allowing for differences in minor allele frequency (MAF) across populations without an increased false positive risk.

The statistical model built into *Tractor* for binary phenotypes tests each SNP for an association with the phenotype using the logistic regression model: logit(Y) = *b*_0_ + *b*_1_*X*_1_ + *b*_2_*X*_2_ + *b*_3_*X*_3_… + *b_k_X_k_* where *X*_1_ is the number of haplotypes of the index ancestry present at that locus for each individual, *X*_2_ is the number of copies of the risk allele coming from the first ancestry, *X*_3_ is the number of copies coming from the second ancestry, and *X*_4_ to *X_k_* are other covariates such as PCs, age, sex, etc. The significance of the risk allele is evaluated with a likelihood ratio test comparing the full model to a model fit without the risk allele, thus allowing estimation of the aggregated effects in the presence of effect size heterogeneity. To further test if a risk allele is ancestry-specific, we evaluate if the difference between *b*_2_ and *b*_3_ is non-zero using a *Z*-test. The model presented is for a 2-way admixed scenario but can be scaled to an arbitrary number of ancestries. We have built pipelines to implement this joint model as well as generate genotype files containing extracted ancestry portions. These tools can be implemented in python and Hail either locally or on the cloud.

## Results

### LAI has high accuracy for African Americans

We ran LAI using the program RFmix, a discriminative approach which estimates local ancestry by using conditional random fields parameterized with random forests^62^. RFmix can run on multi-way admixture populations, outperforms other local ancestry inference methods for minority populations, and leverages the ancestry components in admixed reference panel individuals, highly important when there is a lack of homogenous reference panels - often the case for understudied groups^63,64^. As *Tractor* relies heavily on LAI calls, we first ran simulations to ensure that RFmix called local ancestry accurately. LAI was highly accurate in a realistic demographic model for African American (AA) individuals (one pulse of admixture 9 generations ago with 84% contribution of haplotypes from Africa (AFR) and 16% from Europe (EUR); see *Methods),* assigning the correct ancestry ~98% of the time (Table S1). To ensure that *Tractor* performed well across demographic models, we varied demographic parameters including admixture fractions and pulse timings. Specifically, we varied the pulse of admixture in time to 3 generations and 20 generations ago and changed the admixture fractions to 30/70% and 50/50% EUR and AFR ancestry, respectively (**Figure** S1). We also checked the ancestry-specific accuracy in the realistic demographic scenario to assess if there was a bias in calling dependent on ancestry. Across all demographic models and ancestries, site-wise LAI calls were similarly accurate, with the correct call being obtained ~98% of the time (Table S1). While we refer solely to continental level ancestry here, we appreciate the high level of diversity and admixture within the continents and particularly in Africa. As reference panels for diverse groups grow in size, we will have increased ability to examine more geographically refined groups and deconvolve ever more specific haplotypes.

### Recovery of long-range haplotypes disrupted by statistical phasing

While errors in statistical phasing can lead to errors in LAI, we found that iterating between LAI and statistical phasing improved the accuracy of both. Errors in statistical phasing are a major concern^65,66^, but few methods to recover disrupted haplotypes exist. Taking advantage of the unique ability to visualize tracts offered by admixed individuals, we additionally improve long-range haplotype resolution by correcting chromosome strand switch errors from phasing, which we find to be common in admixed cohorts. We demonstrate that using local ancestry information, we are able to consistently correct switch errors and recover disrupted haplotypes, making tract distributions look significantly more realistic (**Figures** 1,2).

**Figure 1.**
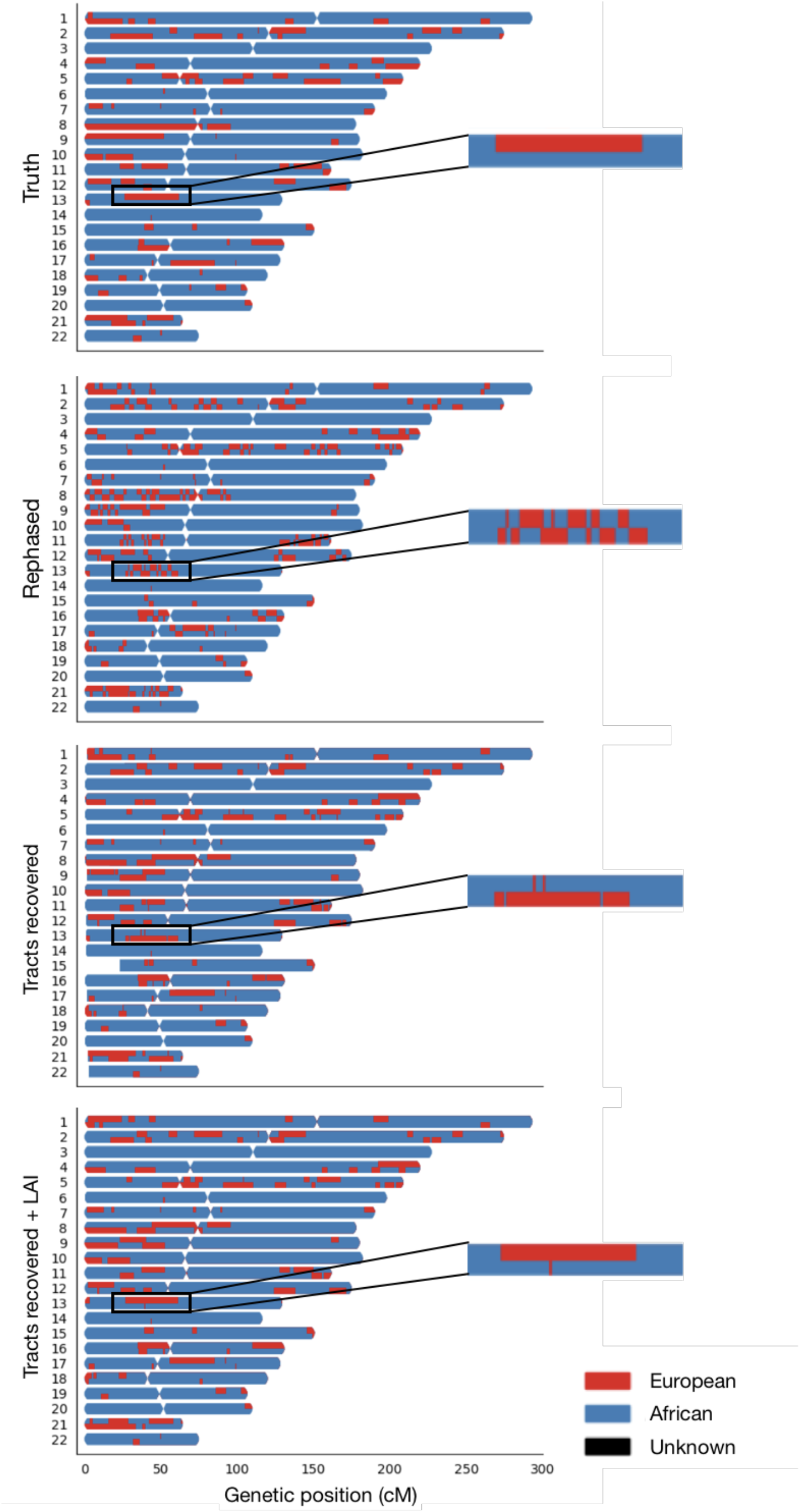
Painted karyograms of a simulated AA individual individual showing EUR (red) and AFR (blue) ancestral tracts across data treatments. The top panel shows the truth results for an example individual in our simulated AA cohort. A painted karyogram after statistical phasing is shown in the second row – note the disruption of long haplotypes. The third panel illustrates our recovery of tracts broken by switch errors in phasing. The bottom panel shows the smoothing and further improvement of tracts acquired through an additional round of LAI. The same section of chr13 showing an example tract at higher resolution is pictured on the right to highlight tract recovery.

**Figure 2.**
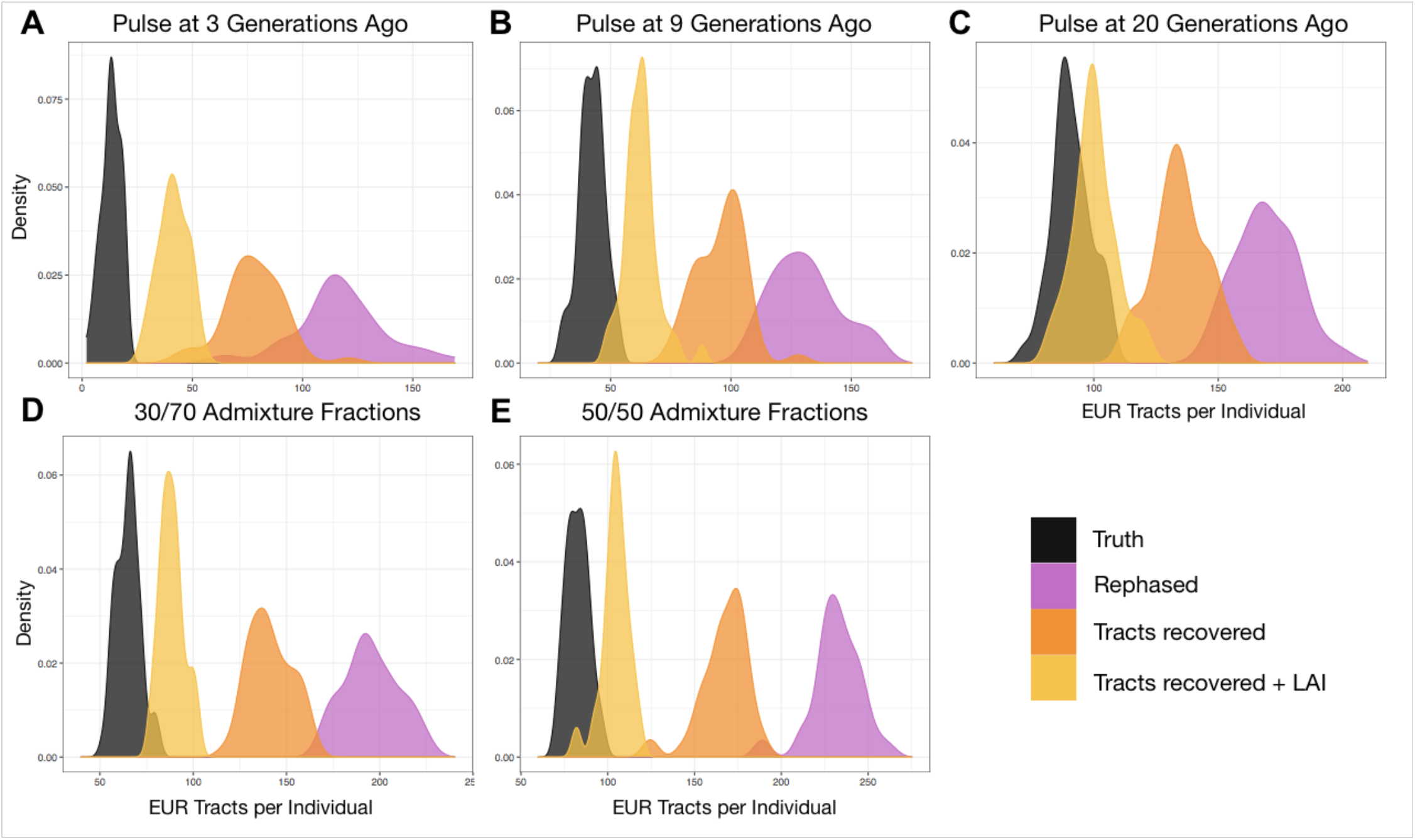
*Tractor* recovers disrupted tracts, improving tract distributions. The top row (**A-C**) shows the improvements to the distributions of the number of discrete EUR tracts observed in simulated AA individuals under demographic models of 1 pulse of admixture at 3, 9 (realistic for AA population history) and 20 generations ago. The bottom row (**D,E**) shows the results from different initial admixture fractions, of 70% and 50% AFR, respectively, at the realistic 9 generations since admixture. These can be compared to the inferred demographic model in AA with ~80% AFR ancestry shown in **B**. In all panels, the simulated truth dataset is shown in black, after statistical phasing in purple, immediately after tract recovery procedures is in orange, and after one additional round of LAI after tract recovery in yellow.

To replicate standard analytical procedures employed on cohort data, we statistically phased our truth dataset using SHAPEIT2^67^ software with a balanced reference panel composed of EUR and AFR continental individuals from the 1000 Genomes Project^39^. We then examined the distribution and lengths of the EUR tracts. Analyzing the less common ancestry tracts allows for more precise quantification of tract counts because it is less likely that recombination will mask their phase switch errors. The probability of obtaining the observed number of tracts after phasing (131 after phasing vs 42 in the truth dataset) given the input demographic model was extremely unlikely, *p*=5.0×10^-26^. After correcting phase switch errors, the likelihood of the tract distribution improved, albeit still with significantly more switches than in the truth data (*p*=2.7×10^-11^, 96 tracts) – approximately half the excess tracts without phase error correction. After then implementing one additional round of LAI on the corrected genotype files, the number of excess tracts was further reduced (*p*=0.009, 62 tracts). Thus, our procedure for correcting phase switch errors successfully recovers long-range haplotypes and better approximates the true tract length distributions (**Figure** 2).

To ensure that phase switch error correction performs well across different population histories, we ran simulations to assess how closely the tract length distributions approximated the truth for a range of demographic models. Specifically, we checked performance varying the timing of admixture pulses (including 3, 9, and 20 generations ago) and the admixture proportions in the simulation (70/30, and 50/50 AFR/EUR, respectively). Under all scenarios, our tract recovery procedure improves strand flips in painted karyograms (**Figures** 1, S1) and decreases the significance of the difference between the observed versus true tract length distributions (**Figure** 2, Table S2). We note that running an additional round of LAI after recovering haplotypes produced the most accurate tract length distributions. This is due to the improved ability of the model to recognize ancestry switch points once more complete haplotypes have been recovered, resulting in smoothing over previous short miscalls.

### Evaluating the landscape of GWAS power gains from Tractor

We simulated individuals’ likelihoods of being cases as a function of AFR admixture fraction, the ancestral haplotype of each copy of the risk allele, and the risk allele dosage (See *Methods, Supplementary Information).* This framework is equivalent to modifying the marginal effect sizes due to a tag SNP for a shared causative mutation being monomorphic in EUR but variable in AFR, which is plausible as individuals from Africa contain almost a million more variants than other populations^68^. This also incorporates the clinically observed phenomenon of disease prevalence differing as a function of ancestry. We then ran association tests and compared the power across the odds ratio spectrum under the traditional GWAS model and under our model. Compared to the traditional model, there is a significant gain in power using the *Tractor* framework with similar improvements across sample sizes and disease prevalences (**Figure** 3). Power increases further when there is a difference in MAF across ancestries.

**Figure 3.**
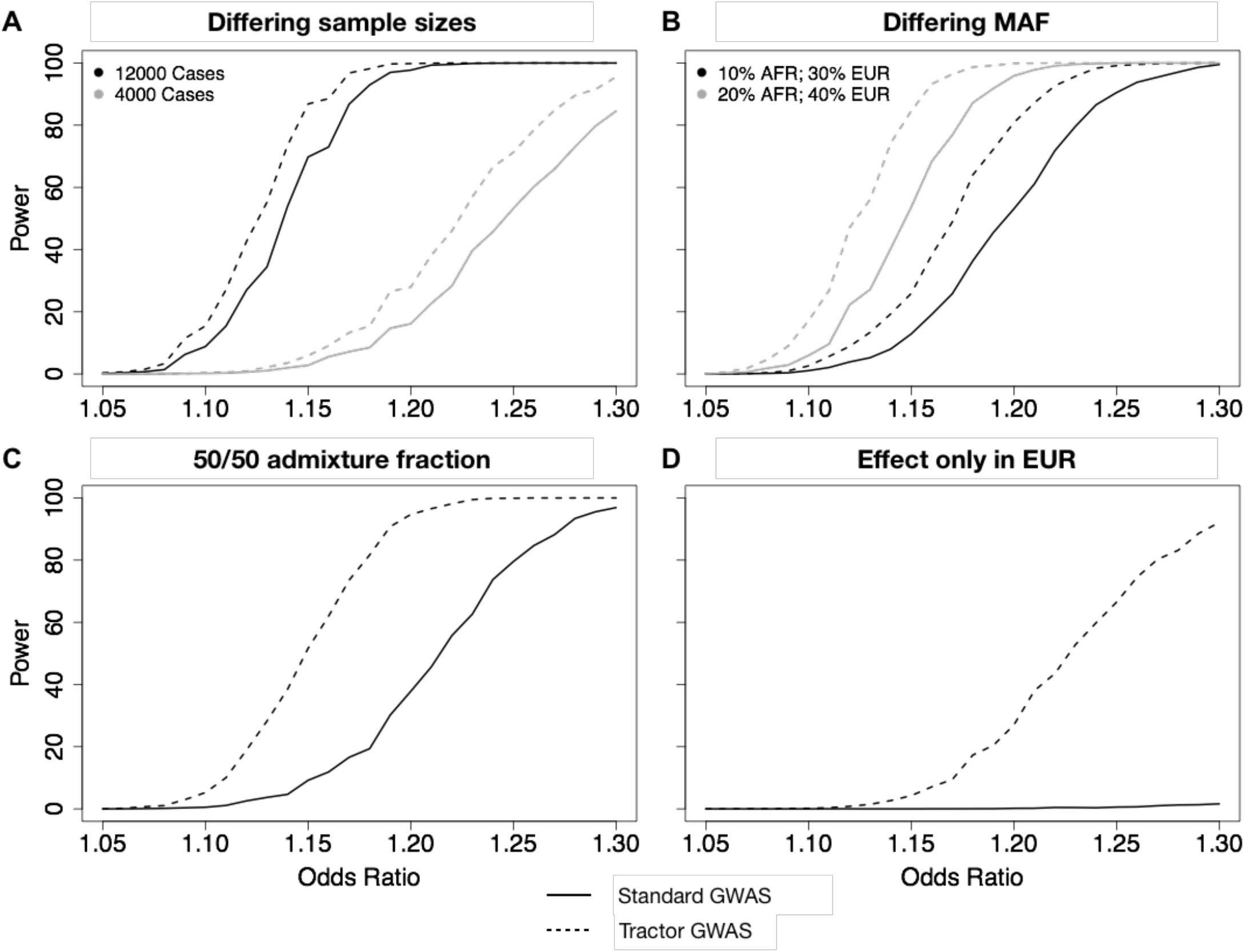
GWAS power gains across sample sizes, ancestral MAF differences, admixture proportions, and effect size differences. In all scenarios shown, dashed lines correspond to the power from the *Tractor* model incorporating local ancestry, solid lines are for a traditional GWAS model. In all panels we modeled a 10% disease prevalence. Unless otherwise noted, we used the parameters for a realistic demographic scenario for AA individuals: 80% AFR ancestry, an effect present only in the AFR genetic background, 12k cases and 30k controls, and 20% MAF. (**A**) There are similar gains in GWAS power when using the *Tractor* LAI-aware model across samples sizes of 4,000 (grey) and 12,000 (black) cases with 2x controls. (**B**) When there is a MAF difference between ancestries, the gains in power are even more pronounced. Gains vary across the allele frequency spectrum: black=MAF 10% AFR, 30% EUR; grey=MAF 20% AFR, 40% EUR. (**C**) Gains become more pronounced when the admixture fractions are modified to 50/50. (**D**) Dramatic gains are obtained when the effect is switched to instead only be present on the EUR background.

We ran similar sets of simulations varying the paraments of the effect size difference, the absolute MAF, MAF difference across ancestries, and admixture fractions (**Figures** 3, S2, S3). The biggest power gain comes if an allelic effect is present in the smaller fraction ancestry only. For example, in a realistic AA demographic model, EUR ancestry makes up only ~20% of the sample. If we model an allele with an effect only active in the EUR background (**Figure** 3D), analyzing the tracts together without LAI information will have essentially no power to detect an association due to the higher noise relative to signal from uninformative tracts. However, *Tractor* is able to recover the effective sample size and power that one would have had if analyzing just the effect haplotypes, i.e. the EUR segments alone.

The scenario where *Tractor* is most powerful is when there is heterogeneity in the apparent effect size for the same variant across ancestries. Such heterogeneity in effect sizes may be a consequence of the same variant having different effects in different populations (e.g., in the context of gene-environment interactions) or may arise from differences in the indirect association evidence of the variant (i.e., the contribution to the estimated effect size from tagging other causal genetic variants). Differences in indirect association can come from ancestry-specific variation or from different patterns of linkage disequilibrium. Moreover, under the converse scenarios where there is predicted to be no benefit, the power loss is minimal—we lose a degree of freedom, resulting in a less precise error estimate for each SNP effect (Figure S2,3). In no case does *Tractor* dramatically underperform compared to the traditional GWAS model. See *Supplementary Information* for additional simulations and detail about power results.

### No increase in false positive rate with the Tractor model

We quantified the false positive rate of the *Tractor* model by simulating a variant with no effect and counting the spurious significant associations identified in a simulated realistic AA population given *α.* = 0.05. Across our tests at various MAFs and between-ancestry MAF differences, we observe no clear difference in false positive rate between *Tractor* and traditional GWAS (**Figure** S4). In addition, we calculated the genomic inflation factor, *λ*_GC_, of null phenotypes across GWAS permutations and confirmed no significant inflation using the *Tractor* GWAS model (**Figure** S5b). Therefore, there does not appear to be an elevation in false positive rates with the *Tractor* framework, suggesting that the observed power increases are from improved detection of true biological signal.

### Tractor replicates known associations and identifies hits for blood lipids in admixed individuals missed by standard GWAS

To ensure that our *Tractor* joint-analysis GWAS model also performs well on empirical data, we ran the method for three well characterized blood lipid phenotypes which have been demonstrated to have ancestry-specific effects: Total Cholesterol (TC), high-density lipoprotein cholesterol (HDL-C), and low-density lipoprotein cholesterol (LDL-C). We constructed a pseudo-cohort of 4309 two-way African-European admixed individuals from the UK Biobank (UKBB) with biomarker phenotype data to serve as our sample (**Figure** S4). *Tractor* GWAS replicated previously implicated associations for blood lipids in this cohort^45,69–71^, reaching the standard genome-wide significance level of 5×10^-8^ at previous top associations, including in genes *PCSK9, LDLR,* and *APOE* (**Figure** 4). In some cases, *Tractor* improved the observed top hit significance.

**Figure 4.**
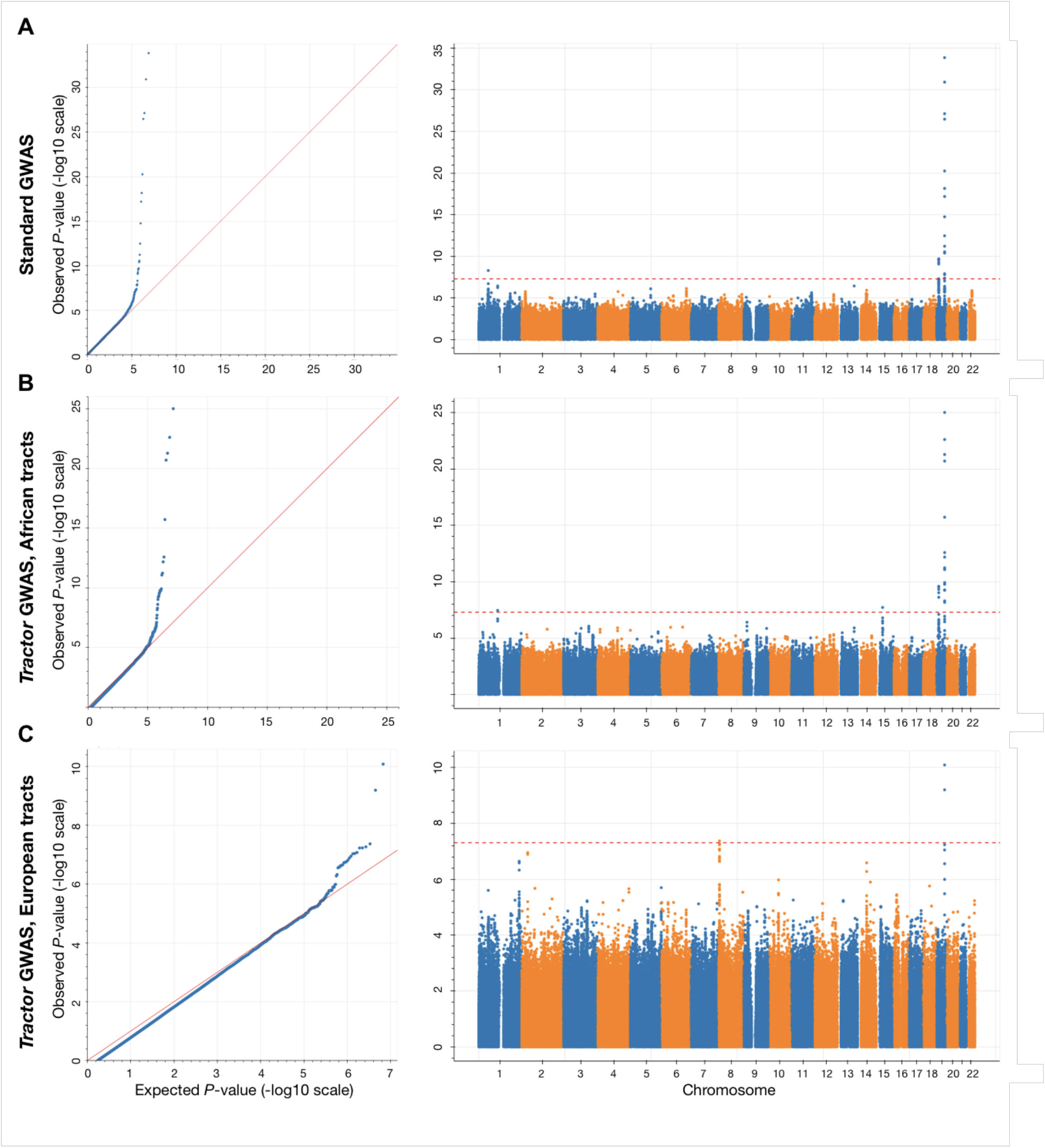
*Tractor* GWAS replicates established hits for Total Cholesterol in admixed African-European individuals and identifies new ancestry-specific loci. QQ and Manhattan plots for Total Cholesterol using the standard GWAS model (**A**) compared to *Tractor* joint-analysis results for the AFR (**B**) and EUR (**C**) backgrounds.

Our LAI-incorporating model was also able to identify hits in these admixed UKBB individuals that standard GWAS was not when using the traditional genome-wide significance threshold (**Figure** 4). For example, we identify a hit missed by standard GWAS on the same dataset that is present only on the AFR background on chromosome 1 (rs12740374, *p*=3.46×10^-8^). This locus has previously been shown to affect blood lipid levels, metabolic syndrome, and coronary heart disease risk in independent AA cohorts^69,72–77^, and was determined to be the causal variant for affecting LDL-C in a multi-ethnic fine-mapping study^78^. Had we not deconvolved ancestral tracts for our GWAS, we would have missed this site with a demonstrated effect on our phenotype and population of interest.

We additionally identify a novel peak on chr15 that only reached significance in the AFR tracts in this UKBB cohort. The lead SNP (rs12594517, p= 1.915×10^-8^) lies in an intergenic area and is uncharacterized. The closest gene neighboring it is *MEIS2,* lying ~70kb upstream, followed by *C15orf41.* While the precise role and mechanism this locus plays in affecting blood lipids remains unclear, *MEIS2* has previously been found to be associated to body mass index and waist circumference and *C15orf41* was a significant hit in a previous GWAS of cholesterol ^79,80^. Though further follow-up is needed to clarify any direct relationship to TC, this association highlights the utility of *Tractor* to identify signals that would be undetectable in admixed cohorts without accounting for local ancestry.

*Tractor* is also able to refine the location of GWAS signals to closer to the causal variant than is possible using standard GWAS procedures. TC has previously been mapped to the gene *DOCK6* in AA cohorts^69^, a finding we replicate for the suggestive GWAS threshold with standard GWAS on UKBB admixed individuals in the same intronic area as previously found. However, when we run the *Tractor* model, we identify a lead *DOCK6* SNP 20kb downstream in the AFR samples, as well as in a meta-analysis of hits from deconvolved AFR and EUR tracts. This new lead SNP (rs2278426) is a missense mutation spanning both *DOCK6* as well as *ANGPTL8,* a gene which may play a key role in blood lipid regulation (see *Discussion,* **Figure** 5). To assess whether our improved ability to localize to this variant was due to a true effect size differences between the EUR and AFR or to a marginal effect size difference driven by MAF or LD differences across the ancestries, we further attempted validating its association by fine-mapping TC in other large-scale populations: 345,235 white British individuals from UKBB and 135,808 Japanese individuals from Biobank Japan (^81^, Ulirsch, JC., Kanai, M. et al., in prep., Kanai, M. et al., in prep). rs2278426 was successfully fine-mapped to a 95% credible set in both populations, with maximum posterior inclusion probability of 0.993 in Biobank Japan. This variant is at 26% frequency in the gnomAD^82^ East Asian ancestry individuals, 18% in African, and 4% frequency in the non-Finish Europeans. These frequency patterns suggest higher power in non-European population to localize a causal variant compared to Europeans. Though below the traditional genome-wide significance level in our sample of ~4300 individuals, this locus highlights the improved ability to localize GWAS signal thanks to leveraging additional breakpoints in admixed genomes.

**Figure 5.**
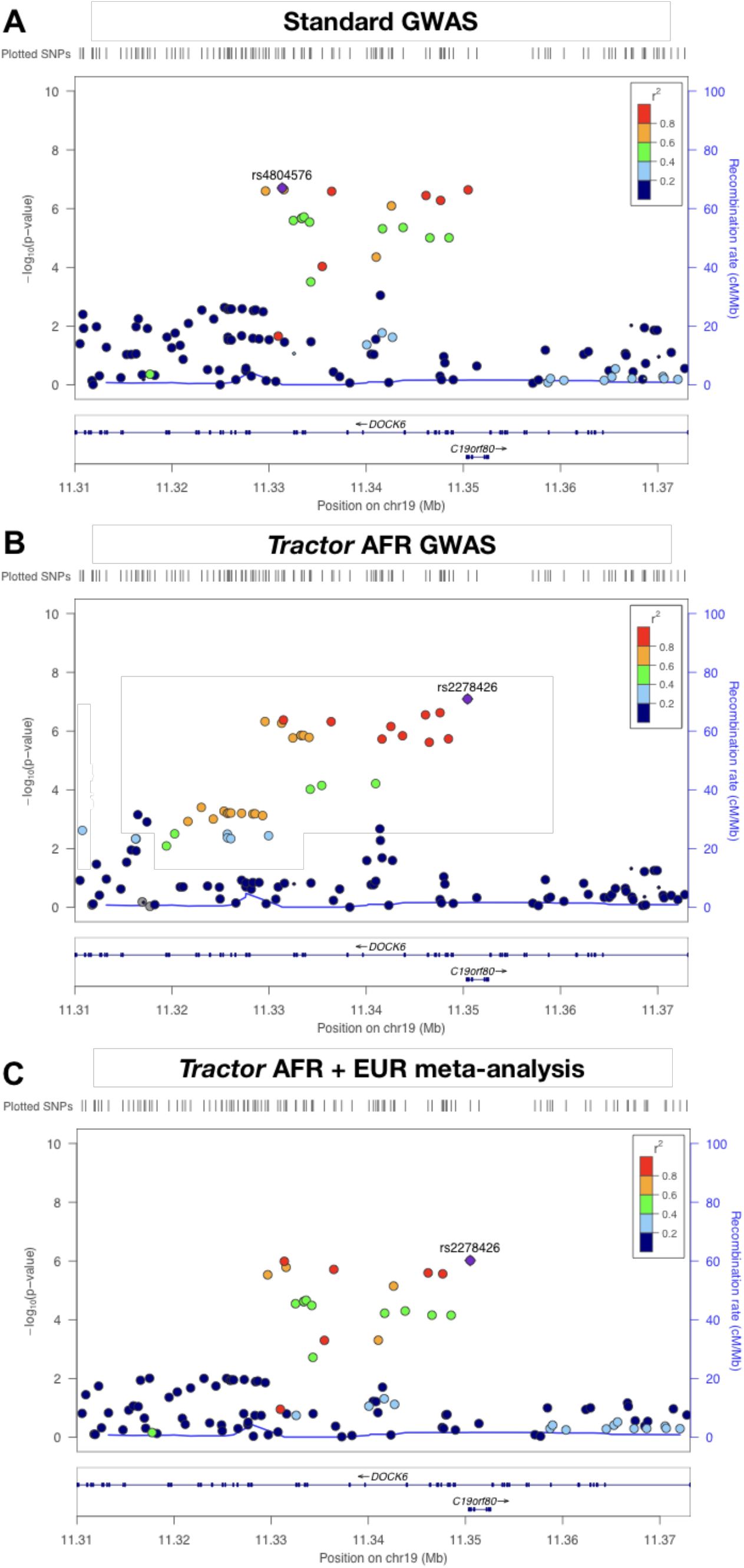
*Tractor* better localizes a top hit for Total Cholesterol. Previous hits for TC had pinpointed *DOCK6* as the gene of interest. Comparing runs on UKBB admixed individuals with a standard GWAS model (**A**), AFR-specific GWAS with *Tractor* (**B**), and a meta-analysis of GWAS runs on deconvolved EUR and AFR tracts (**C**), both ancestry-specific runs pinpoint a lead SNP ~20kb downstream in an intron of *DOCK6* spanning a better candidate gene *ANGPTL8* (also known as C19orf80) as the lead SNP. No significant signal was seen in the EUR segments. In all plots, point size is proportional to the number of samples included for that test, and color indicates *r*^2^ to the named lead SNP. For **B**, the recombination rate line was generated from the AFR superpopulation of the 1000 Genomes Project, for other panels the EUR superpopulation rate is shown.

## Discussion

Despite the recent advances in understanding the genetics of complex diseases, major limitations remain in our knowledge of the architecture of such disorders in minority and admixed populations. Here, we present an analytical framework and statistical gene discovery method distributed as a scalable software package named *Tractor,* which allows admixed samples to be appropriately included alongside homogenous ones in a well calibrated manner in statistical genomics efforts. We test our framework in a simulation model designed to emulate real AA cohorts. We also apply it to empirical data from admixed African-descent individuals of the UKBB. We observe a gain in power to detect risk loci across sample sizes, demographic models, and disease prevalences using the *Tractor* framework, particularly when effect sizes are heterogeneous across populations. Our approach incorporates a local-ancestry aware GWAS method that can extend the traditional GWAS model. *Tractor* generates ancestry specific *p* values and effect size estimates, which admixture mapping cannot, and which can be extremely helpful in post-GWAS efforts such as constructing genetic risk scores for understudied populations. We demonstrate that our framework also gives increased precision in localizing GWAS signal by leveraging the disrupted LD blocks visible with ancestral chromosome painting in recently admixed groups. This reduces the credible set of SNPs and aids in the prioritization of variants for subsequent functional testing.

The *Tractor* pipeline requires several inputs, most importantly accurate local ancestry calls. Users should ensure good LAI performance in their target cohorts. A major determinant of accurate LAI calls is a comprehensive and well-matched reference panel^49,83^. Of relevance is that reference panels are more plentiful for Eurasian populations than for other groups, underscoring the need to expand sequencing efforts in more global populations. To ensure LAI was unbiased across regions of the genome in our GWAS, we examined the distribution of local ancestry across the genome. Local ancestry inference appears relatively evenly distributed across the genome and proportional to global admixture proportions (**Figures** 1, S6). However, we caution that calls around centromeres and at the ends of chromosomes are most likely to include error, as these genomic regions do not have anchor points on one edge. We similarly recommend that LAI ideally to be conducted on whole genome sequencing data to avoid the introduction of biases. We also note that we have thoroughly tested *Tractor* here in the two-way admixture model that reflects AA demographic history. The analytic infrastructure, however, can currently also run on a three-way admixed model and our statistical model can scale to an arbitrary number of ancestries. Future work will test power and optimize the code in a variety of multi-way admixed demographic scenarios. A final consideration is to ensure consistent phenotyping across ancestry groups, as is standard in multi-ethnic GWAS.

We thoroughly evaluated the landscape of when *Tractor* does and does not add power to association studies in simulated data modeled after AA cohorts of the PGC-PTSD (**Figures** 3, S2, S3). In situations where there are differences across ancestries, *Tractor* recovers the power that would be lost from analyzing populations together. In particular, power gains are greatest when there is an effect size difference at a locus between ancestries coupled with differing MAF. For example, our simulated case of a 20% MAF difference for an allele with an effect only in the AFR genetic background would allow for identification of risk variants with an odds ratio ~0.1 smaller than would be possible with traditional GWAS. This allows for detection of additional loci that would have been undetected without modeling local ancestry. *Tractor* can also boost power when there is an effect only on one haplotype background or an allele only present in one ancestry, again most dramatically when that ancestry is less frequent in the dataset. In such an instance in a standard GWAS setting, the signal would be dramatically reduced due to noise from the uninformative majority haplotypes, resulting in extremely low power to detect the locus (**Figure** 3D). Power can be recovered, however, by deconvolving local ancestry and analyzing genotypes on ancestry-specific haplotypes, thus controlling for population structure as well as identifying risk variants that would otherwise be undetectable.

Conversely, we find that it is not generally necessary to include local ancestry in a GWAS model when there is no effect size difference between groups. Notably, we are referring here to detection of marginal effect sizes as well as true effect sizes. There is evidence suggesting that in most cases (with some notable exceptions^8,69,84^), the true effect sizes of causative variants are likely to be equal across ancestries^33,38,85–90^. However, the marginal effect size of a tag SNP might routinely be different across ancestries due to differences in ancestral MAF and the LD patterns resulting from each ancestral population’s demographic history^91,92^. We underscore that power gains appear to be from true biological signal rather than false positives, as we quantified the *Tractor* false positive rate to be no higher than standard GWAS (Figures S4, S5).

We would like to highlight that *Tractor* benefits from power gains to detect the marginal beta (pertaining to, for example, an allele which is only present in one population and tags a nearby causal variant), in addition to the rarer case of variants with true effects only on one haplotypic backbone. Our framework therefore will improve power at substantially more locations across the genome than only at sites which have ancestry-specific causal effect differences. *Tractor* also benefits from increased power in cases where functionally important (and likely rare) alleles only present in one population are missed by genotyping or imputation. In such situations the common alleles in LD, despite being shared across populations, would be associated to the phenotype as a function of which haplotypic background they are found on, and thus would have a haplotype-specific effect. Another relevant scenario to consider would be the presence of LD in regions where there are ancestry-specific markers intermingled with shared ones. This would affect the univariate scan results such that considering the haplotypic background on which alleles fall would particularly aid in localizing signal through improved marginal beta estimates, even when the causal effect is the same in both ancestries.

*Tractor* was also able to replicate established GWAS hits, discover new ones, and aid in the localization of GWAS signal with empirical data. We replicated known hits for well-characterized blood lipid phenotypes when testing *Tractor* on a dataset consisting of ~4300 2-way admixed African-European individuals from the UKBB (**Figure** 4). We further demonstrate an improved ability to localize GWAS signal to putative causal SNPs previously identified in another diverse collection, Biobank Japan^81^ (**Figure** 5). Specifically, previous analyses of blood lipid phenotypes in an admixed AA cohort had pinpointed the TC top hit to lie within the gene *DOCK6*^69^, nearby the lead SNP for this region in standard GWAS on the admixed UKBB individuals (rs4804576). The *Tractor* AFR ancestry, as well as results from a meta-analyses of summary statistics from AFR and EUR deconvolved genotype files, identified a different top association ~20kb downstream (rs2278426) which additionally lies over the *ANGPTL8* gene on the positive strand while spanning an intronic area of *DOCK6* on the minus strand. *ANGPTL8,* also known as lipasin and betatrophin, has been shown to regulate plasma lipid levels in mice by inhibiting the enzyme lipoprotein lipase^93–96^. In humans, ANGPTL8 levels correlate with metabolic phenotypes including type 2 diabetes and obesity^97–100^ and HDL-C expression levels across diverse ancestry groups have been demonstrated to better correlate with than *DOCK6*^101^. Together these make *ANGPTL8* a more promising candidate gene than *DOCK6,* which has no clear tie to blood lipid phenotypes. Intriguingly, the *Tractor* lead SNP, rs2278426, is a missense mutation in *ANGPTL8* (p.Arg59Trp) that is predicted to be possibly damaging and deleterious by polyphen and SIFT, respectively^102,103^. This variant is at 18% frequency in gnomAD^82^ AFR individuals but at 4% frequency in the non-Finnish Europeans. These frequency differences highlight how leveraging different diverse populations allows for the improved identification of risk variants as well as how employing multi-ethnic mapping methods aids in the resolution of association signals.

The *Tractor* infrastructure released here may be helpful in multiple statistical genetics use cases beyond GWAS. For example, correcting for population structure should be a key early step in evolutionary genomic studies on admixed populations running analyses such as genome-wide scans of selection to avoid bias in selection statistics^104–106^. Within medical genetics, accounting for the ancestral background on which an allele appears will be key in admixed populations, particularly in studies of rarer variants which are more population specific^107,108^. For example, because AFR and EUR haplotypes have different rates of background variation^68^, controlling for the local ancestral background may help pinpoint the differences between cases and controls in burden testing that would previously have been overwhelmed by uninformative markers.

In sum, *Tractor* allows users to account for genotype-level ancestry in a precise manner, allowing for the well-calibrated inclusion of admixed individuals in large-scale gene discovery efforts. This approach provides a number of benefits over traditional GWAS, including the production of ancestry-specific effect size estimates a *p* values, improved localization of GWAS signal, and power boosts in genetic scenarios such as when there are effect size or MAF differences across ancestries. This infrastructure is designed as a series of steps to be flexible and easily ported into other statistical genomics activities. We freely provide *Tractor* code in python and Hail^109^, a scalable cloud-compatible framework, as well as examples of implementation in a Jupyter notebook^110^. *Tractor* advances the existing methodologies for studying the genetics of complex disorders in admixed populations.

## Online Methods

### QC and LAI Pipeline

The core feature of the *Tractor* framework relies on accounting for fine-scale population structure as informed by local ancestry (i.e. ancestral chromosome painting). *Tractor* then uses this information to (i) correct for individuals’ ancestral dosage at all variant sites, (ii) recover long-range tracts in admixed individuals; and (iii) extract the tracts and ancestry dosage counts from each ancestry component for use in ancestryspecific association tests. We have tested and built this framework around LAI calls from RFmix versions 1 and 2^62^, and have built an automated pipeline (https://github.com/eatkinson/Post-QC) to perform all necessary post-genotyping QC, data harmonization, phasing, and LA inference to consistently prepare the data for downstream analysis. The main code is in bash, subscripts are written in python (See *Supplementary Information* for additional details).

In all tests shown here, we ran RFmix_v2 with 1 EM iteration and a window size of 0.2 cM. We used the HapMap b37 recombination map^111^ to inform switch locations. The -n 5 flag (terminal node size for random forest trees) was included to account for an unequal number of reference individuals per reference population. We additionally used the --reanalyze-reference flag, which recalculates admixture in the reference samples themselves for improved ability to distinguish ancestries. This is especially important when the reference samples are themselves admixed. As a reference panel for our 2-way admixed simulated African-European cohorts, we used relevant populations of the 1000G reference panel given *a priori* knowledge of AA’s demographic history^112–114^ consisting of 108 YRI and 99 CEU. Painted karyogram plots were produced using a modified version of publicly available code (https://github.com/armartin/ancestry_pipeline). We have optimized this pipeline under the two-way admixed AA demographic scenario. *Tractor* additionally supports 3-way admixture calls with an expanded set of scripts (also at https://github.com/eatkinson/Tractor). In all cases we recommend conducting tests of LAI accuracy to ensure reliability, as accurate LAI calls are required for good performance.

### LAI Accuracy

We validated that LAI was performing well in the AA use case. To do this, we generated a truth dataset by simulating individuals with known phase and LA from empirical data. Our simulation reference panel consisted of haplotypes from homogenous PGC-PTSD individuals who had ≥95% EUR or AFR ancestry as inferred by SNPweights^115^. We simulated admixture between these reference individuals with admix-simu^116^ using a realistic demographic scenario for the AA population^113,114^ of 1 pulse of admixture 9 generations ago with 84% contribution from Africa and 16% from Europe. The resultant population mixes amongst itself until the present day, copying haplotypes from the previous generation with break points informed by the HapMap combined recombination map^111^. This retains the LD structure and genetic variation present in real genomic data and ensures that the truth dataset resembles cohort data as closely as possible. We then called LA with the 1000 Genomes^68^ AFR and EUR superpopulations as our reference panel, and calculated LAI accuracy as how often the ancestry call was correct in the simulated truth data.

### Correcting Switch Errors from Statistical Phasing using Local Ancestry

Despite LAI calling ancestry dosage accurately, frequent chromosomal switches were visible in painted karyograms (**Figure** 1), which we determined were due to phasing errors. It is important to retain complete tracts, as spurious breakpoints will reduce the accuracy haplotype-based test. *Tractor* detects and fixes phase switches using the most likely ancestry assignment of subpopulations as determined by a conditional random field from RFmix. We define phase switches as a swap of ancestry across a chromosome within a 1 cM window at a region with heterozygous ancestry dosage.

To ensure that correcting phase switch errors improved results compared to the truth expectations for the input demographic scenario, we modeled the expected distributions of EUR tract lengths within AA individuals using a Poisson process with rate=9, the number of generations ago when the pulse of admixture occurred (**Figure** 2). The waiting time until a recombination event disrupts a tract is expected to follow this distribution, with a slight shortening of tracts proportional to the percent admixture due to the inability to visualize tract switches that occur across regions of the same ancestry. The overall proportion of the genome in the realistic scenarios was within range of expectations given the simulation model of 16% European, 84% African ancestry (15.1% and 84.9%, respectively).

### GWAS power simulations incorporating local ancestry

We assessed the improvements in GWAS power from using *Tractor* through simulations. We formulated our simulation framework on the suggestions of Skotte et al (2019)^117^. Power calculations were based on a simulation framework that initially models an AA population assuming a bi-allelic disease risk allele with a 20% overall MAF and an additive effect in the AFR genetic background but not in the EUR. Specifically, the overall admixture proportions were drawn from a beta distribution with shape parameters 7.76 and 2.17, the fitted parameters to this distribution for AFR ancestry proportions observed in the PGC-PTSD Freeze 2 AA cohorts. The genotype of each copy of the allele was drawn from a binomial distribution with the probability of having the minor allele set to the MAF. We simulated a disease phenotype with individuals’ risk drawn from a binomial distribution assuming a 10% disease prevalence. Risk of developing the phenotype was modified on a log-additive scale according to the admixture proportions and the presence of the minor allele on an AFR background using a logit model. In this model, the probability of disease was set to −2.19 + log of allelic risk effect size*number of copies of the minor allele coming an AFR ancestral background + 0.5*AFR admixture proportion. −2.19 was chosen as it represents a 10% probability of disease given no AFR admixture or copies of the minor allele from either ancestral background. The 0.5 value in 0.5*AFR Admixture was set in order to induce stratification in the simulated population, as is observed in empirical data. In other words, all of our simulations modeled increasing disease prevalence with admixture fractions, reflective of clinical observation. With this simulation design, individuals with higher AFR ancestry proportions are more likely to be cases whereas those with higher EUR ancestry proportions are more likely to be controls. Subjects’ disease status was then drawn from a binomial distribution with the probability parameterized to their individual disease risk according to the logit model. Cases and controls were sampled at random from the simulated population at a 2.5:1 control to case ratio, the approximate ratio of controls to cases in PGC-PTSD freeze 2.

Under each simulation, we fit three logistic regression models of disease status that included: M1) admixture only, M2) number of copies of the risk allele only, and M3) admixture + number of copies of the risk allele on a EUR background + number of copies on an AFR background. M1 serves as a null comparison to evaluate the significance of including the SNP as a predictor. The significance of M2 and M3 are evaluated by likelihood ratio tests comparing them to M1. For each 100 simulations at a given effect size and sample size, for both M2 and M3 we estimated power as the proportion of the time that the likelihood ratio test was significant (p < 5e-8). We performed 100 rounds of simulation with this model at each level of allelic effect size ranging from Odds Ratio (OR) 1.05 to 1.3 and case sample size *N*=4000 and 12000.

### Characterizing the landscape of Tractor power across genomic and disease contexts

To evaluate *Tractor* power gains, we ran similar sets of simulations varying effect size differences across ancestries, MAF differences, admixture fractions, and disease prevalence (**Figures** 3, S2, S3).

#### Varying effect size across populations

To examine the effect of modifying the effect sizes, we introduced an effect on EUR haplotypes as well, rather than just on AFR. All these simulations assumed 80% admixture, 10% disease prevalence, and 20% MAF in both groups. We modeled cases across the OR spectrum where there was an effect of equal size in both ancestries, a 30% larger effect size on the EUR background, an effect size 30% larger on the AFR haplotype, and an effect size only in the EUR.

#### Varying absolute MAF

We next fixed all other parameters and modified the absolute MAF of the simulated risk allele, with the relative difference in MAF between ancestries remaining constant. We changed our MAF from 20% to 10% and 40% under both the models of an effect only in the AFR background and with matching effect sizes between EUR and AFR.

#### MAF differences between groups

To see if having a difference in the MAF between the two ancestral groups affected GWAS power, we varied the MAF in the EUR background to be 10, 20, and 30% while keeping the AFR MAF set to 20%.

#### False positive rate

We quantified the false positive rate by simulating a variant with no effect and counting significant associations identified in a simulated realistic AA population given *α.* = 0.05.

### Selection of two-way admixed African-European empirical individuals

To select individuals with 2-way admixture with European and West African ancestry, we took a two-pronged approach. First, we combined genetic reference data from the 1000 Genomes Project^39^ and Human Genome Diversity Panel^118^, then harmonized meta-data according to consistent continental ancestries. We then ran PCA on unrelated individuals from the reference dataset. To partition individuals in the UKBB based on their continental ancestry, we used the PC loadings from the reference dataset to project UK Biobank individuals into the same PC space. We trained a random forest classifier given continental ancestry meta-data based on the top 6 PCs from the reference training data. We applied this random forest to the projected UK Biobank PCA data and assigned AFR ancestries if the random forest probability was >50%, otherwise individuals were dropped from further analysis.

For those individuals classified by their genetic data to have AFR ancestry, we then combined the 1000 Genomes and Human Genome Diversity Panel reference data with genetic data from the African Genome Variation Project as well as these UKBB individuals. To restrict to only two-way admixed West African-European ancestry individuals, we restricted to individuals with at least 12.5% European ancestry, at least 10% African ancestry, and who did not deviate more than 1 standard deviation from the AFR-EUR cline (**Figure** S6A, B). This resulted in approximately 4300 individuals per blood lipid trait. Global ancestry fraction estimates were obtained from running ADMIXTURE^119^ with *k*=2 (which was the best fit *k* value to this dataset based on 5-fold cross-validation) on these individuals with 1000 Genomes Project^39^ EUR and AFR superpopulation individuals as reference data (**Figure** S6C). To ensure there were no major areas of the genome where local ancestry inference was skewing significantly from the expected global fractions, we also assessed the cumulative local ancestry calls across the genome for the UKBB admixed subset (**Figure** S6D).

### Software implementations

We developed separate scripts to deconvolve ancestry tracts and calculate haplotype dosages, correct phase switch errors, and run a *Tractor* GWAS to obtain ancestry-specific effect size estimates and *p* values. Pre-GWAS steps are available as independent python scripts. We separated steps to allow for maximum flexibility when using *Tractor.* To implement the joint modeling GWAS approach with the novel linear regression model described here, we have built a scalable pipeline in Hail^109^ which can be implemented locally or on the Google Cloud Platform^120^. Descriptions of the steps and an example Jupyter notebook^110^ demonstrating analytical steps and visualization of results of the *Tractor* joint-analysis GWAS are freely available on github (https://github.com/eatkinson/Tractor).

An alternative pipeline designed for use across environments where Hail may not be as readily implemented involves running the separate/meta-analysis GWAS version of *Tractor.* This pipeline requires the initial processing steps to optionally correct phase switch errors and deconvolve ancestry tracts into their own VCF files. Next, GWAS can be run for the deconvolved files containing different ancestral components with the user’s preferred GWAS software, such as plink^121^. In this implementation, a standard GWAS model can be run on each ancestral component separately using the ancestry-specific VCF output by *Tractor,* which contains fully or partially missing data including only haplotypes from the ancestry in question. Results from the different ancestry runs could then be meta-analyzed to increase power by incorporating summary statistics from both populations, though we recommend preferentially using the joint-analysis method described in this manuscript to avoid any potential bias from combining multiple ancestral portions of the genome of the same individuals. This implementation is also compatible in large-scale collections where there are large numbers of homogenous individuals, for example many Europeans, but too limited a number of admixed individuals to be run in a GWAS alone. The EUR sections of the admixed cohorts could be analyzed alongside the homogenous European cohorts, making better use of the admixed samples even if other ancestry portions are not utilized, and increasing the effective sample size.

### Empirical test of Tractor on blood lipid phenotypes in European-African admixed UKBB individuals

To ensure that *Tractor* replicated well-established associations, we ran standard GWAS, the *Tractor* joint-analysis model, and a meta-analysis of summary statistics from EUR and AFR deconvolved tracts on ~4300 admixed African-European individuals from the UKBB on the biomarker blood lipid traits of Total Cholesterol (TC), high-density lipoprotein cholesterol (HDLC), and low-density lipoprotein cholesterol (LDLC). We included covariates capturing global ancestry, age, sex, and blood dilution factor in all runs. We assessed meta-analysis performance using different metrics to capture global ancestry, namely the first 20 PCs versus the AFR fraction as determined by ADMIXTURE, which did not have substantive differences. In the jointanalysis framework, we used the measure of global AFR ancestry fraction to more directly capture global ancestry and avoid any potential collinearity with local ancestry from PCs. We generated QQ plots alongside each trait and compared the inflation of test statistics in each GWAS case by looking at the genomic inflation factor, *λ*_GC_. We then compared results to those obtained from the same individuals using a standard GWAS approach. No individuals overlap between the previous study of interest (Natarajan et al. 2018, which includes diverse TOPMed^24^ individuals) and the UKBB individuals included here. As expected, near-identical results were obtained from the meta- and joint-approaches. Gene visualizations were produced with LocusZoom^122^, Manhattan and QQ plots with bokeh^123^.

### Statistical fine-mapping of top hits in independent cohorts

We conducted GWAS and statistical fine-mapping in two additional large-scale cohorts, 345,235 white British individuals from UKBB and 135,808 Japanese individuals from BBJ. For the UKBB white British, we used previously conducted fine-mapping results for TC (https://www.finucanelab.org/data). Briefly, we computed association statistics for the variants with INFO > 0.8, MAF > 0.01 % (except for rare coding variants with MAC > 0), and HWE p-value > 1e-10 using BOLT-LMM^124^ with covariates including the top 20 PCs, sex, age, age^2^, sex * age, sex * age^2^, and blood dilution factor. We used FINEMAP v1.3.1^125,126^ and susieR v0.8.1.0521^127^ for fine-mapping using the GWAS summary statistics and in-sample dosage LD matrices computed by LDstore v2.0b. We defined regions based on 3 Mb window surrounding lead variants and merged them if overlapped. The maximum number of causal variants in a region was specified as 10. For BBJ, we additionally conducted fine-mapping using the same pipeline as we did for UKBB. The GWAS summary statistics of TC was computed for the variants with Rsq > 0.7 and MAF > 0.01% using BOLT-LMM with the covariates including top 20 PCs, sex, age, age^2^, sex * age, sex * age^2^, and disease status (affected versus non-affected) for the 47 target diseases in the BBJ. The details about genotyping and imputation was extensively described previously^81,128^.

### Assessment of the correct empirical p value for admixed individuals

To evaluate the appropriate *p* value threshold for *Tractor* associations, we estimated ancestry-specific empirical null *p* value distributions via permutation. Although the genome-wide significance threshold (*p* < 5 × 10^-8^) is widely adopted in the current literature, previous work has shown that different ancestry groups have different numbers of independent variants^68^. Here, we permuted a null continuous phenotype 1,000 times using the same admixed African-European individuals from UKBB as in the *Tractor* cholesterol GWAS to assess the correct *p* value threshold for the admixed individuals in this study. We measured the minimum *p* values of associations (*p*_min_) for each ancestry and derived an ancestry-specific empirical genome-wide significance threshold as the fifth percentile *(α* = 0.05) of *p*min across permutations as previously described^129^.

We calculated this percentile using the Harrell–Davis distribution-free quantile estimator^130^ and calculated the 95% confidence interval via bootstrapping. Based on the permutation results (**Figure** S5a), we defined a studywide significance threshold at a conservative level of *p* = 1 × 10^-8^ for both AFR- and EUR-specific associations. In addition, we calculated the genomic inflation factor, *λ*_GC_, of null phenotypes across permutations and confirmed no significant inflation using the *Tractor* GWAS model (**Figure** S5b).

## Acknowledgements

We thank Pradeep Natarajan, Sarah Gagliano Taliun, and many other scientists within and beyond Boston for their intellectual contributions to this work. This project was supported by the National Institute of Mental Health (K01 MH121659 and T32 MH017119 to E.G.A.; K99MH117229 to A.R.M.; 2R01MH106595 to C.M.N. and K.C. K.). M.K. was supported by a Nakajima Foundation Fellowship and the Masason Foundation. M.L.S. was supported by Fundacao de Amparo a Pesquisa do Estado de Sao Paulo (#2018/09328-2). The BioBank Japan Project was supported by the Tailor-Made Medical Treatment Program of the Ministry of Education, Culture, Sports, Science, and Technology (MEXT), the Japan Agency for Medical Research and Development (AMED).

## Author Contributions

E.G.A. designed and implemented the pipeline, ran analyses, and drafted the primary manuscript. A.X.M. designed and ran analyses. M.K. designed and ran analyses with the aid of J.C.U., Y.K., Y.O., and H.K.F. A.R.M. contributed code and aided in writing the manuscript. K.J.K. and M.S. aided in code implementation.

K.C.K, C.M. N., B.M.N. and M.J.D. supervised and advised on the project. All authors reviewed and approved the final draft.

## Code availability

All code is freely available. *Tractor* scripts, in both cloud executable Hail format and python formats, can be found at https://github.com/eatkinson/Tractor. The automated QC pipeline to prepare datasets for *Tractor* and run LAI is located at https://github.com/eatkinson/Post-QC.

