## Supplementary Information for "*Tractor*: A framework allowing for improved inclusion of admixed individuals in large-scale association studies"

***Automated Genotype QC pipeline***

We have additionally built a pipeline that can be used to filter data as a precursor to running *Tractor* (https://github.com/eatkinson/Post-QC). To run this optional precursor step to clean post-QC genotype data before LAI, the user will simply need to input paths to their binary plink format sample genotypes file and a reference panel on the command line. The pipeline will perform all necessary post-genotyping QC, intersecting, phasing, and LAI to consistently prepare the data for downstream analysis. The Post-QC steps that are implement are: (1) Extract only autosomes in data file; (2) Find and remove duplicate loci; (3) Update SNP IDs to dbsnp 144[1]; (4) Orient data to 1000 genome reference[2]; (5) Remove ambiguous A/T and G/C loci. This involves three substeps at step 4: a. Find and remove indels; b. Find and remove loci not found in 1000 genome, or that have different coding alleles than 1000 genome (tri-allelic, for example); c. Flip alleles that are on the wrong strand. This script also outputs warnings if more than 2% of sites are incongruous between the input dataset and 1000G locations, as this can indicate genome build discrepancies. A/T and G/C loci are unable to be strand-resolved and for this reason are routinely removed in QC pipelines. Post-QCed cohort data is then intersected and jointly phased with a user-specified reference panel of individuals. The merged dataset is then filtered to include only informative SNPs present in both the cohort data and reference panel using a MAF filter of 0.5% and a genotype missingness cutoff of 90%. The program Shapeit2[3] is implemented for phasing with switches informed by the HapMap combined recombination map[4].

***Characterizing the landscape of Tractor power across genomic and disease contexts***

To evaluate *Tractor* power gains, we ran similar sets of simulations as described in the main text, varying effect size differences across ancestries, MAF differences, admixture fractions, and disease prevalence (**Figure**s 3,S2, S3). Additional discussion on these simulation results is presented here to provide further detail on those not discussed in the main text.

*Varying absolute MAF:* We next fixed all other parameters and modified the absolute MAF of the simulated risk allele, with the relative difference in MAF between ancestries remaining constant. We changed our MAF from 20% to 10% and 40% under both the models of an effect only in the AFR background and with matching effect sizes between EUR and AFR haplotype context. We note that different MAFs affect power as they would under a regular GWAS; i.e. power tends to increase with MAF (**Figure** S2).

*MAF differences between groups*: To see if having a difference in the MAF between the two ancestral groups affected GWAS power, we varied the MAF in the EUR background to be 10, 20, and 30% while keeping the AFR MAF set to 20%. Changing the MAF differences when the effect sizes are equal across ancestries does not dramatically affect power (Figure S3). Given unequal effect sizes, however, when the effect is present only in the AFR, as the MAF decreases in EUR, the power gain also decreases (**Figure** S3). And vice versa, higher MAF in the non-effect group affords our method more power gains (**Figure** S3).

*Varying admixture proportions*: Admixture proportion affects power in our method because it determines the effective number of samples per ancestry group. Consider the scenario again where there is no effect in EUR but there is an effect in AFR. A populations with 80% AFR ancestry has a greater number of haplotypes in which there is an effect compared to a population with 50% AFR ancestry. As a consequence, the population with 50% AFR admixture has lower power because there are fewer AFR tracts in general, meaning 50% of the sample does not contribute signal as opposed to only 20% of the sample not contributing signal (**Figure** 3).

A related observation of note is that under the assumption of AFR only allelic effects, the simulated power of *Tractor* tended to be quite close to power predicted at *N**average African ancestry proportion using Quanto[5]. This implies that the power of *Tractor* under these conditions is effectively the power of an analysis stratified to just AFR ancestry individuals; i.e. the effective *N* given the AFR ancestral tracts.

*Varying the simulated disease prevalence:* A comparison of two simulated diseases with prevalences of 10% and 20% shows that the relative differences in power between populations remains similar. Because this was a case/control sampling design with no misclassification of disease status, the prevalence of the disease did not notably impact power. In other words, prevalence affects power in our model as it would under a traditional GWAS framework. A theoretical situation in which admixture influenced misdiagnosis of a disease may affect power.

*False positive rate*: We quantified the false positive rate by simulating a variant with no effect and counting spurious significant associations in a simulated realistic AA population given $\boldsymbol{\alpha} = 0.05$. Across our tests at various MAFs and MAF differences at large sample size, we observe no clear difference in false positive rate between *Tractor* and traditional GWAS. Multiple runs of 100 simulations produced uncorrelated results, suggesting any variation in rates is not systematic (**Figure** S3). Therefore, there does not appear to be an elevation in false positive rates with the *Tractor* framework, suggesting that power increases are from detection of true biological signal.


**Supplementary Tables**

**Table S1: LAI accuracy across demographic contexts.** LAI accuracy, defined as the proportion of the time the ancestry of a variant was correctly called in a simulated truth dataset, was calculated for all sites across a range of demographic scenarios modifying the timing of the pulse of admixture in time or the overall fraction of ancestry contribution from the two ancestries. We also calculated ancestry-specific accuracy under the realistic demographic scenario of 1 pulse of admixture 9 generations ago, with 84% contribution from Africa and 16% from Europe. Modified admixture fractions show the proportion of AFR and EUR ancestries, respectively.

| Demographic model | Accuracy |
|---|---|
| *Realistic, global* | 0.979 |
| *Realistic, African* | 0.978 |
| *Realistic, European* | 0.984 |
| *Pulse at 3 gens ago* | 0.984 |
| *Pulse at 20 gens ago* | 0.977 |
| *Admixture of 30/70* | 0.979 |
| *Admixture of 50/50* | 0.982 |

**Table S2. Tract distributions across data treatments and demographic models.** Results shown are for the numbers of lower frequency EUR tracts in simulated AA individuals. *P* values indicate the significance of obtaining the observed number of tracts given the truth expectations a modeled with a Poisson process.

| | ***Truth*** | ***Rephased*** | | ***Unkinked*** | | ***Iterated*** | |
|---|---|---|---|---|---|---|---|
| *Demographic model* | *# Tracts* | *p* | *# Tracts* | *p* | *# Tracts* | *p* | *# Tracts* |
| *Pulse at 3 gens ago* | 13.16 | 2.931E-67 | 117.74 | 8.188E-34 | 78.34 | 1.252E-09 | 41.40 |
| *Pulse at 5 gens ago* | 21.76 | 9.547E-49 | 121.36 | 3.137E-24 | 85.52 | 2.286E-06 | 48.52 |
| *Pulse at 9 gens ago* | 42.22 | 4.997E-26 | 130.94 | 2.752E-11 | 95.84 | 9.039E-03 | 62.62 |
| *Pulse at 15 gens ago* | 69.92 | 2.369E-15 | 149.80 | 2.543E-06 | 116.72 | 8.106E-02 | 84.46 |
| *Pulse at 20 gens ago* | 91.06 | 0.000E+00 | 169.16 | 4.026E-05 | 134.46 | 3.121E-02 | 100.04 |
| *Admixture of 30/70* | 65.10 | 0.000E+00 | 195.78 | 6.034E-16 | 141.34 | 2.291E-02 | 88.72 |
| *Admixture of 50/50* | 82.50 | 0.000E+00 | 231.46 | 0.000E+00 | 167.04 | 2.242E-02 | 104.46 |

### Supplementary Figures

**Figure S1. Painted karyograms of a simulated AFR individual showing EUR (red) and AFR (blue) ancestral tracts across demographic scenarios.** The first column shows the results for the demographic model of one pulse of admixture 3 generations ago, the middle column shows the realistic African American demographic model of one pulse 9 generations ago, and the right column shows a pulse 20 generations ago. In all cases the model involved 84% AFR ancestry and 16% EUR. The rows show the results from treatments of the data across steps. The top row shows the truth results from our simulations. Painted karyograms after statistical phasing of this truth cohort is shown in the second row. The third row illustrates the recovery of tracts broken by switch errors in phasing obtained by unkinking. The bottom row shows the smoothing and further improvement of tracts acquired through an additional round of LAI.

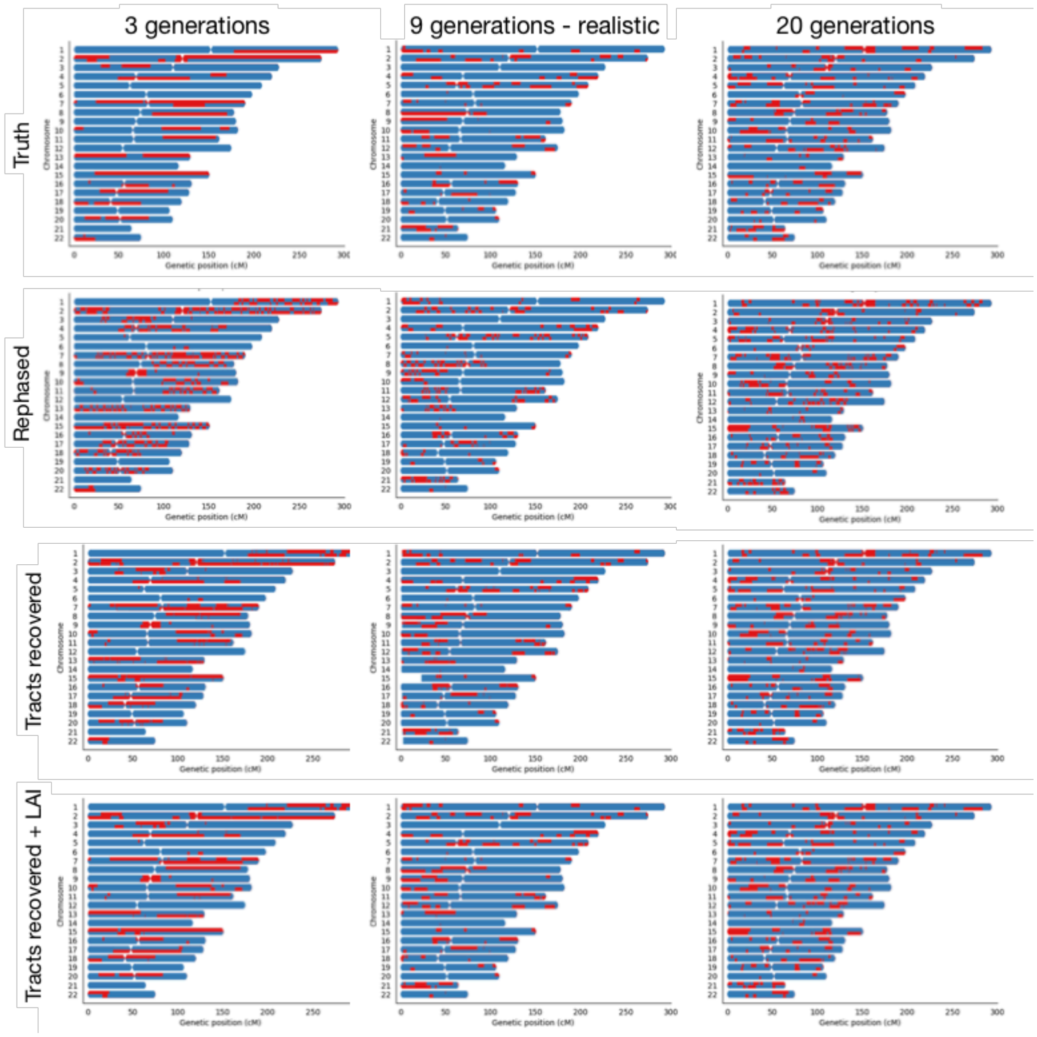

**Figure S2. The contribution of absolute MAF and effect size to *Tractor* power.** All cases assume 80% admixture and 10% disease prevalence with an effect only in the AFR genetic background for 12,000 cases and 30,000 controls. In each panel, the grey solid line uses a traditional GWAS model while the black dashed line is our LAI-incorporating model. Top (**A,B**): Matching effect size between EUR and AFR with shifted absolute MAF. Bottom (**C,D**): effect only in AFR background. Left (**A,C**): MAF is set to 10% in both AFR and EUR. Right (**B,D**): MAF is set to 40% in both AFR and EUR.

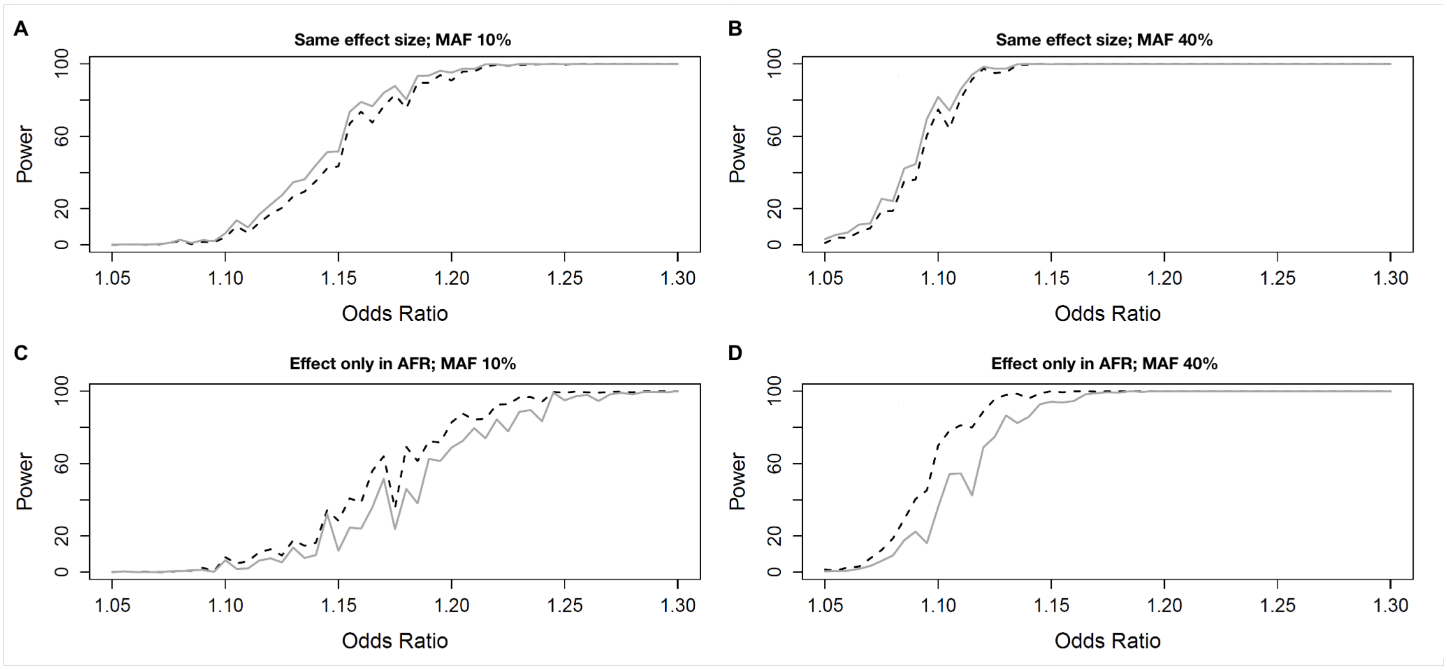


**Figure S3. The interaction of between-ancestry MAF differences and effect sizes on *Tractor* power.** In all cases, the grey solid line uses a traditional GWAS model while the black dashed line is our LAI-incorporating model, admixture proportions are 80% African, disease prevalence is 10%, and the AFR MAF is fixed at 20%. **A,C,E** model the same effect size between EUR and AFR while varying the EUR MAF. **B,D,F** model the case when there is no effect in the EUR background while varying EUR MAF. **G** models an effect size difference of 30% with the effect being stronger in the EUR background at 20% EUR MAF, **H** an effect size difference of 30% with the effect being stronger in the AFR background at 20% EUR MAF.

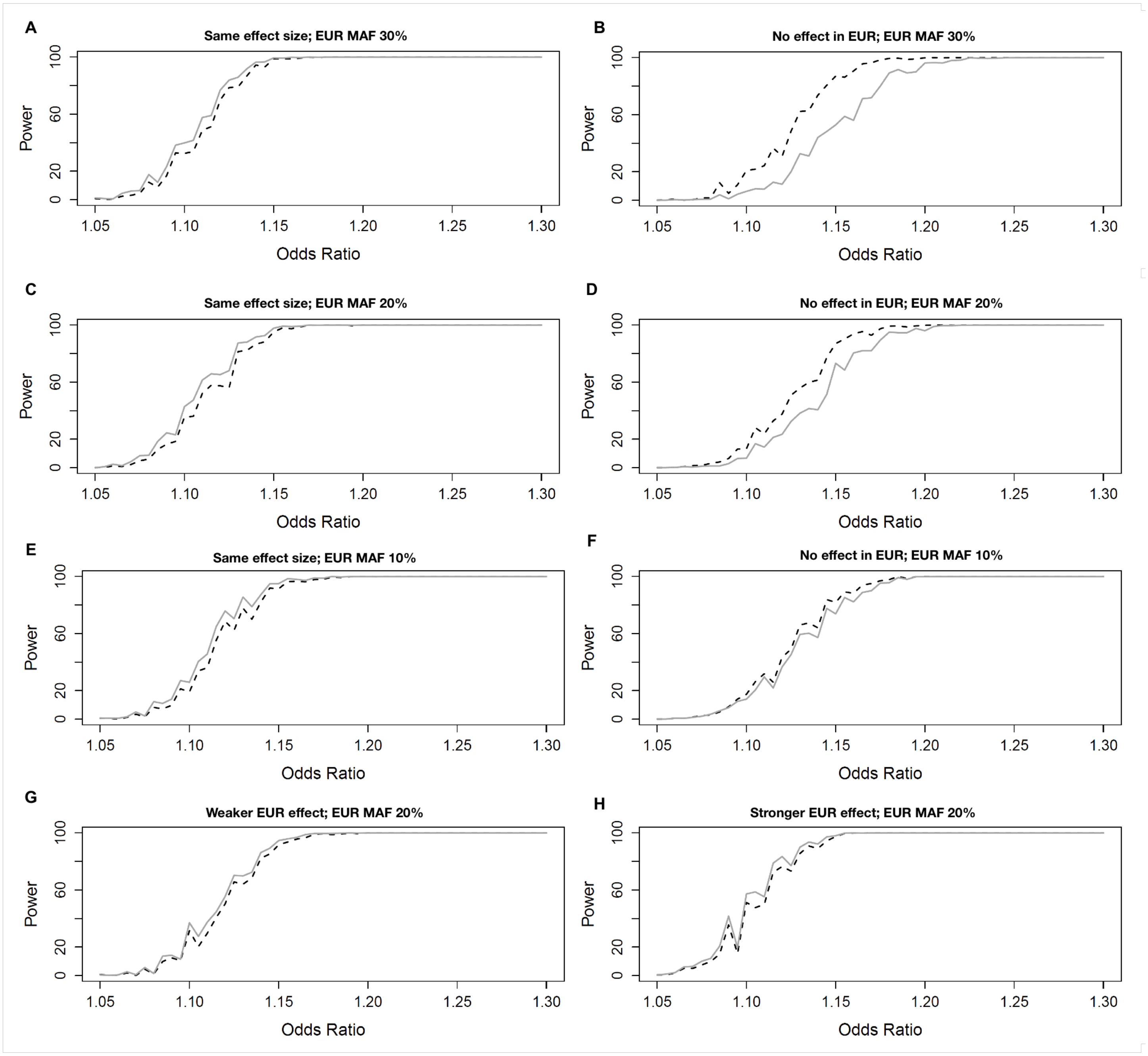

**Figure S4**. **False positive rate of Tractor versus standard GWAS at differing MAFs**. The false positive rate was quantified as the frequency at **α** =0.05 where an allele simulated to have no effect was detected at genome-wide significance. No consistent difference in false positive rates was observed comparing standard and *Tractor* GWAS.

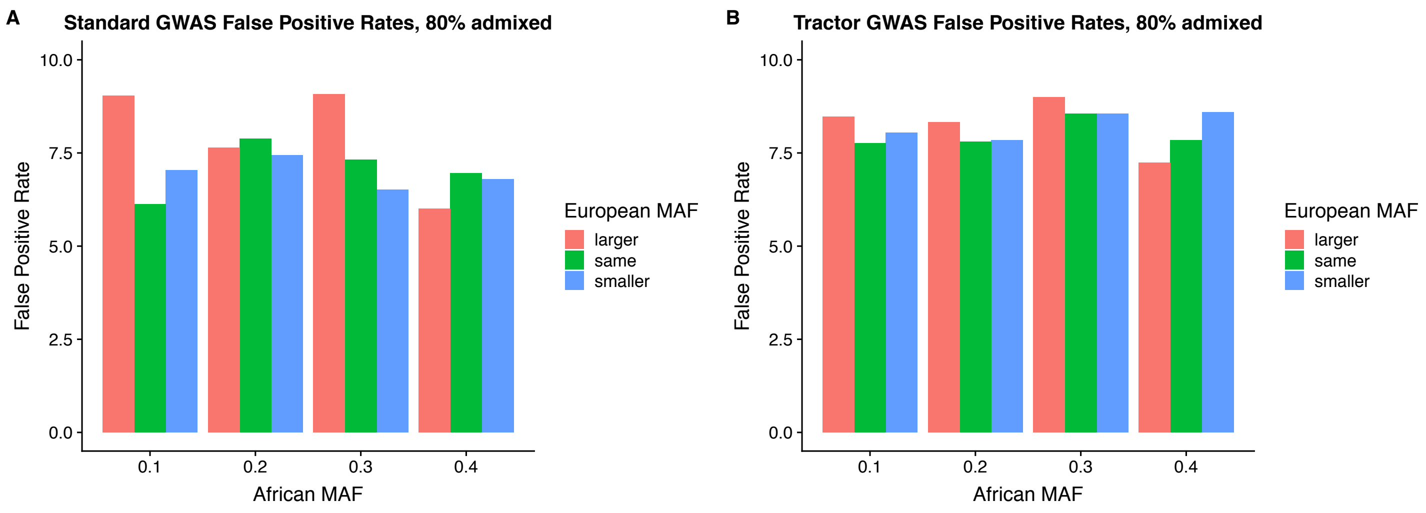


**Figure S5**. Empirical assessment of the ideal genome-wide *p* value threshold for ancestral segments deconvolved by *Tractor*. (a) A slightly more stringent threshold of $p = 1 \times 10^{-8}$ for both AFR- and EUR-specific associations was deemed appropriate for admixed African-European cohorts, consistent with the presence of additional recombination breakpoints present in such admixed populations. (B) No inflation was observed using the *Tractor* GWAS model in permutation tests.

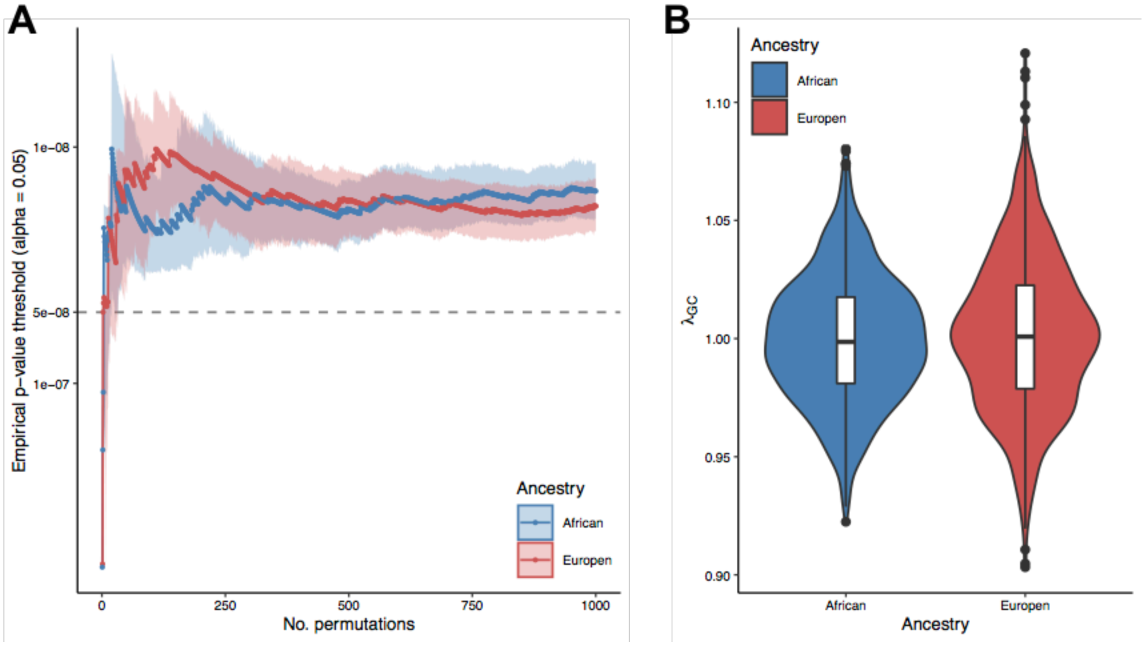


**Figure S6: Selection of admixed African-European UKBB individuals.** (**A**) Ancestry assignment by a random forest model trained on 1000G data. (**B**) Individuals selected for inclusion in two-way admixed subset. (**C**) Admixture fractions at *k*=2 across UKBB individuals included in analyses. The 1000 Genomes EUR and AFR superpopulation individuals were used as a reference for ADMIXTURE. (**D**) Relatively even average distribution of local ancestry across the genome in UKBB admixed individuals. Some edges of chromosomes were trimmed due to uncertainty in calls. Consistent with coloring scheme throughout paper, blue is African ancestry, red is European ancestry.

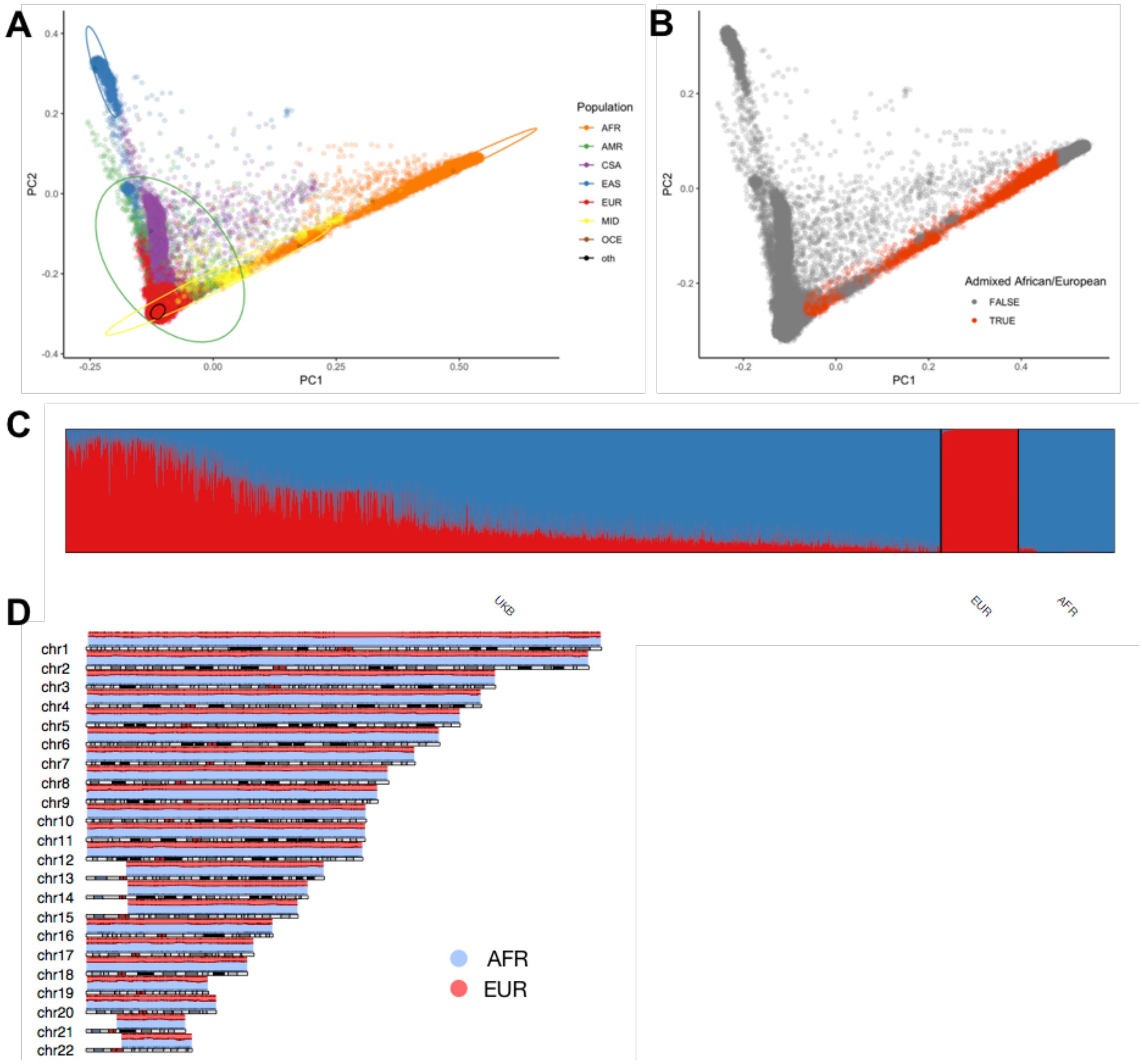
A
B
C
D
Population
AFR
AMR
CSA
EAS
EUR
MID
OCE
oth
PC1
PC2
Admixed African/European
FALSE
TRUE
UKB
EUR
AFR
chr1
chr2
chr3
chr4
chr5
chr6
chr7
chr8
chr9
chr10
chr11
chr12
chr13
chr14
chr15
chr16
chr17
chr18
chr19
chr20
chr21
chr22
AFR
EUR
